# Mutagenesis Induced by Protonation of Single-Stranded DNA is Linked to Glycolytic Sugar Metabolism

**DOI:** 10.1101/2022.07.14.500049

**Authors:** Suzana P. Gelova, Kin Chan

## Abstract

Mutagenesis can be thought of as random, in the sense that the occurrence of each mutational event cannot be predicted with precision in space or time. However, when sufficiently large numbers of mutations are analyzed, recurrent patterns of base changes called mutational signatures can be identified. To date, some 60 single base substitution or SBS signatures have been derived from analysis of cancer genomics data. We recently reported that the ubiquitous signature SBS5 matches the pattern of single nucleotide polymorphisms (SNPs) in humans and has analogs in many species. Using a temperature-sensitive single-stranded DNA mutation reporter system, we also showed that a similar mutational pattern in yeast is dependent on error-prone translesion DNA synthesis and glycolytic sugar metabolism. Here, we further investigated mechanisms that are responsible for this form of mutagenesis in yeast. We first confirmed that excess sugar metabolism leads to increased mutation rate, which was detectable by fluctuation assay. We then ruled out a significant role for aerobic respiration in SBS5-like mutagenesis by observing that petite and wild-type cells did not exhibit statistical differences in mutation frequencies. Since glycolysis is known to produce excess protons, we then investigated the effects of experimental manipulations on pH and mutagenesis. We hypothesized that yeast metabolizing 8% glucose would produce more excess protons than cells metabolizing 2% glucose. Consistent with this, cells metabolizing 8% glucose had lower intracellular and extracellular pH values. Similarly, deletion of *vma3* (encoding a vacuolar H^+^-ATPase subunit) increased mutagenesis. We also found that treating cells with edelfosine (which renders membranes more permeable, including to protons) or culturing in low pH media increased mutagenesis. Altogether, our results agree with multiple biochemical studies showing that protonation of nitrogenous bases can alter base pairing so as to stabilize some mispairs, and shed new light on a common form of intrinsic mutagenesis.

**Graphical Abstract:** 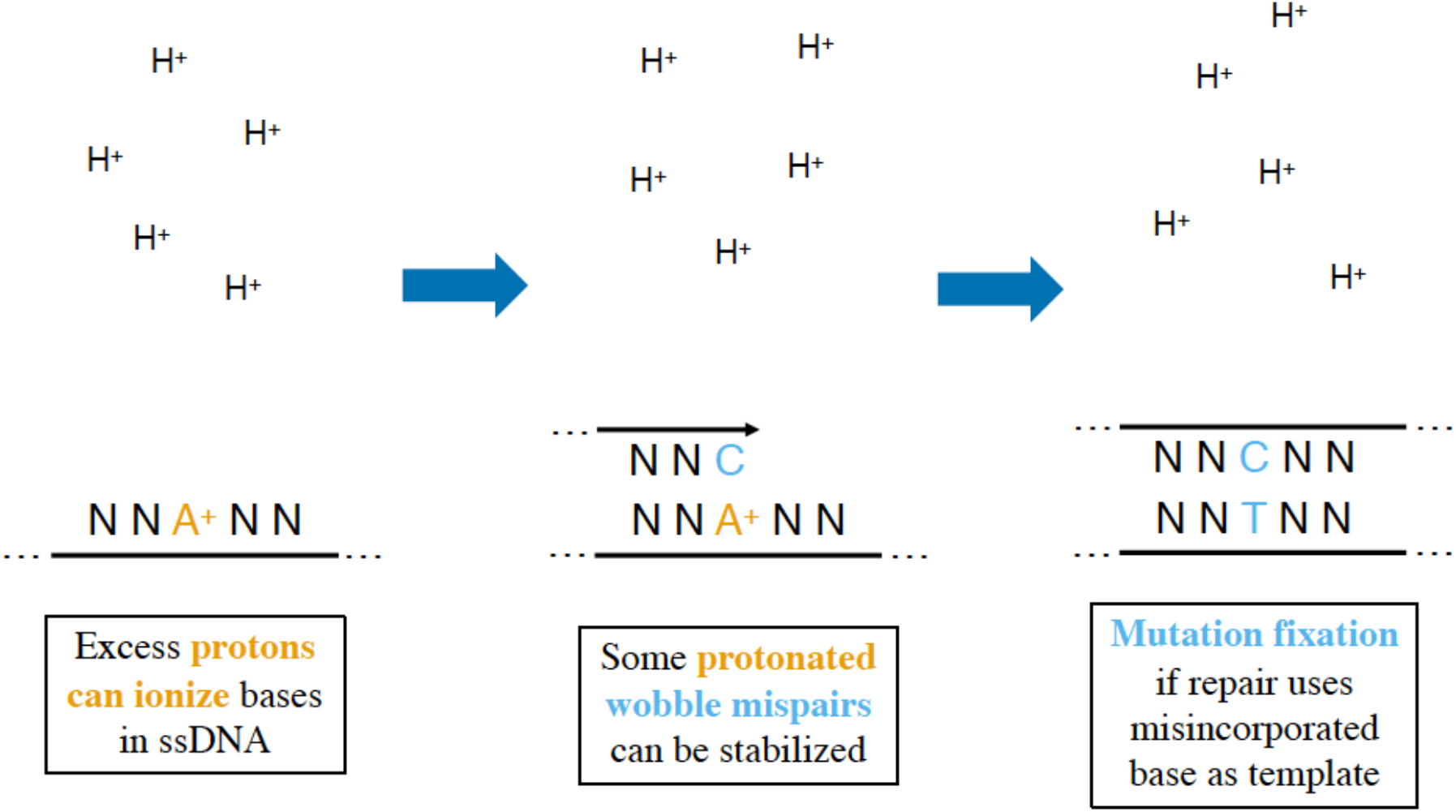

## Introduction

Mutations are the starting material for evolution by natural selection. Mutations oftentimes result after DNA damage from either endogenous processes [1] or exogenous exposures [2–4]. Endogenous DNA damaging processes include the following: deamination of cytosine or 5-methylcytosine to uracil or thymine, respectively [5–9]; base oxidation [10,11]; base alkylation [5,12,13]; glycosidic bond breakage [5,14–16]; single- and double-stranded DNA breaks [5,17,18]. Exogenous DNA damage can take on many forms, including: aflatoxin [19]; aristolochic acid [20]; ionizing radiation [4]; tobacco [21]; and ultraviolet light [3]. Spontaneous ionization or isomerization (i.e., tautomerization) of DNA bases can also alter base pairing characteristics, and are thought to be another possible source of mutations [22–25].

It is important to note that these and other DNA damaging processes or exposures do not affect all bases equally. Adenine, cytosine, guanine, and thymine in DNA each have their own specific set of reactive chemical groups (i.e., amines, carbonyls, labile ring atoms) [26]. The local sequence context can also be a key factor affecting susceptibility to DNA damage. As a result, any given DNA damaging process or agent may be more likely to react with some base(s) than others, yielding recurrent, reproducible patterns of sequence changes that have been termed mutational signatures [27].

Mutational signatures typically are extracted from genomic data sets using the non-negative matrix factorization (NMF) algorithm [28]. There are approximately 60 single base substitution (SBS) signatures described in the <u>C</u>atalog <u>o</u>f <u>S</u>omatic <u>M</u>utations <u>i</u>n <u>C</u>ancer (or COSMIC) at this point [29]. Many mutational signatures have well-defined etiologies, such as: SBS1 (i.e., <u>S</u>ingle <u>B</u>ase <u>S</u>ubstitution signature 1) from deamination of 5-methylcytosine at <u>C</u>pG motifs; SBS2 and SBS13 from enzymatic deamination of cytosine at TC motifs by APOBEC enzymes; SBS3 from deficiencies in homologous recombination DNA repair; SBS4 and SBS29 from tobacco smoking and chewing, respectively; SBS6, SBS15, SBS21, SBS26, and SBS44 from various deficiencies in DNA mismatch repair; SBS7a through SBS7d from ultraviolet light exposure; SBS10a through SBS10d from mutations of the replicative DNA polymerases epsilon or delta; SBS18 from reactive oxygen species; SBS30 and SBS36 from DNA base excision repair deficiencies [27]. Currently, about one-third of the previously described mutational signatures remain of undefined etiology [29]. Further, with more sequencing data being produced on a continuing basis, more mutational signatures surely remain to be discovered as well.

In spite of the great deal of progress made to elucidate the etiologies of mutational signatures, one of the most common remains relatively poorly understood. This signature, SBS5, is found in all cancer types and is present in almost all individual cancer samples, being typically one of the most prominent signatures in each sample [27]. The numbers of mutations attributed to SBS5 increase steadily as we age; interestingly, there is considerable variation in the extent of this “clock-like” mutation accumulation effect between individuals, i.e., some have many SBS5 mutations while others have relatively few [30]. Furthermore, over 70% of mutations in early human embryos [31] and ~75% of de novo mutations in human children [32] fit the SBS5 pattern. In contrast to other mutational signatures which can diminish in prevalence as cancers undergo subclonal diversification, SBS5 usually remains a prominent signature that contributes significantly to intratumor heterogeneity [33].

Previously, we showed that SBS5 is the human counterpart in a continuum of similar intrinsic mutational patterns in many species. Indeed, the pattern of human single nucleotide polymorphisms (SNPs) also matches SBS5 very closely [34]. Additionally, we showed that a similar mutational pattern can be recapitulated using a baker’s yeast (*Saccharomyces cerevisiae*) reporter system featuring controlled generation of genomic single-stranded DNA (ssDNA) [35]. The SBS5-like mutational pattern is dependent on error-prone translesion DNA synthesis in regions of ssDNA and sugar metabolism. More SBS5-like mutagenesis is observed when *cdc13-1* cells are arrested at the G2 phase of the cell cycle in media with more glucose. Further, we showed that blocking the completion of glycolysis using an auxin-inducible degron allele of the principle pyruvate kinase gene abrogates ~90% of the SBS5-like mutagenesis [34].

In this work, we investigated mechanisms contributing to this form of mutagenesis further. Our results show that excess sugar metabolism, in addition to increasing mutagenesis, has important effects on lowering both intracellular and extracellular pH. Moreover, our findings implicate protonation of ssDNA as a likely source of the increased mutagenesis in our highly sensitive mutation reporter system. Our study provides evidence that ionization of exposed nitrogenous bases in ssDNA within intact living cells likely alters base pairing characteristics, resulting in higher likelihood of mutation.

## Materials and Methods

### Reagents and Consumables

Bacto peptone (product code 211677) and yeast extract (212750) were purchased from Becton, Dickinson and Co. (Franklin Lakes, NJ). Canavanine (C9758), adenine sulfate dihydrate (AD0028), and digitonin (D141) were purchased from MilliporeSigma (St. Louis, MO). Agar (FB0010), glucose (GB0219), hygromycin (BS725), PCR purification spin column kit (BS654), plasmid DNA spin column kit (BS614), agarose (D0012), Tris-Borate-EDTA (TBE) buffer (A0026), ethidium bromide (D0197), sorbitol (SB0491), KH2PO4 (PBR0445), Na_2_HPO_4_ (S0404), NaCl (SB0476), 96-Well 2 mL Deep Plates (BR581-96NS), and amino acids or nucleotides (besides adenine) for media preparation were purchased from BioBasic (Markham, ON). KCl was purchased from Merck (Darmstadt, Germany). G418 sulfate (450-130) was purchased from Wisent (St-Bruno, QC). Q5 high fidelity DNA polymerase PCR kit (M0491), T4 DNA ligase (M0202), XhoI (R0146), and XbaI (R0145) were purchased from New England Biolabs Canada (Whitby, ON). Microseals ‘B’ were purchased from Bio-Rad (Hercules, CA). CELLSTAR black polystyrene clear bottom 96 well microtiter plates were purchased from Greiner Bio-One (Monroe, NC). Phosphate buffered saline (PBS) was prepared as a 10× stock (pH 7.4) with the following components at these final concentrations: KH2PO4 (18 mM), Na_2_HPO_4_ (100 mM), NaCl (1.37 M), and KCl (27 mM). Plasmid p426MET25_sfpHluorin (MRV55) was purchased from Addgene (Watertown, MA). DH5α *E. coli* competent cells (18265-017) were purchased from Invitrogen (Waltham, MA).

### Yeast Media

Routine culturing was done on YPDA rich solid media (2% Bacto peptone, 1% Bacto yeast extract, 2% glucose, supplemented with 0.001% adenine sulfate, 2% agar) or YPDA liquid rich media (2% Bacto peptone, 1% Bacto yeast extract, 2% glucose, supplemented with 0.001% adenine sulfate), as appropriate. Liquid rich media with 8% glucose (YPD8A) were same as baseline YPDA, except containing 8% glucose. Similarly, for YPDA with 6% sorbitol. Percentage values all refer to weight by volume (w/v).

For synthetic complete solid media, 1 L consisted of: 20 g agar, 20 g D-glucose, 5 g ammonium sulfate, 1.7 g yeast nitrogen base without amino acids or ammonium sulfate, 60 mg adenine sulfate, 50 mg L-arginine HCl, 75 mg L-aspartic acid, 100 mg L-glutamic acid, 20 mg L-histidine HCl, 50 mg L-isoleucine, 100 mg L-leucine, 120 mg L-lysine HCl, 20 mg L-methionine, 50 mg L-phenylalanine, 375 mg L-serine, 100 mg L-threonine, 50 mg L-tryptophan, 50 mg L-tyrosine, 150 mg L-valine, 60 mg uracil, and deionized water to 1 L total volume. pH was adjusted to 5.8 before autoclaving.

For canavanine media with low (0.33 ×) adenine, 1 L consisted of: 20 g agar, 20 g D-glucose, 5 g ammonium sulfate, 1.7 g yeast nitrogen base without amino acids or ammonium sulfate, 20 mg adenine sulfate, 75 mg L-aspartic acid, 100 mg L-glutamic acid, 20 mg L-histidine HCl, 50 mg L-isoleucine, 100 mg L-leucine, 120 mg L-lysine HCl, 20 mg L-methionine, 50 mg L-phenylalanine, 375 mg L-serine, 100 mg L-threonine, 50 mg L-tryptophan, 50 mg L-tyrosine, 150 mg L-valine, 60 mg uracil, and deionized water to 1 L total volume. pH was adjusted to 5.8 before autoclaving. After cooling media to ≤ 60°C, 6 mL of 1% canavanine sulfate solution (filter sterilized) was added and stirred thoroughly to mix, before pouring plates.

### Yeast Genetics

The ySR127 yeast strain (a *MATα* haploid bearing the *cdc13-1* temperature sensitive allele [36,37]) and its derivatives were used in this study. ySR127 is itself derived from strain CG379 [38] and has a cassette of three reporter genes (*CAN1, URA3*, and *ADE2*) near the de novo left telomere of chromosome V. These three genes had been deleted from their native loci. Additional details about ySR127 were described previously [35]. All strains and plasmids are listed in Table 1, and are available upon request.

**Table 1:**
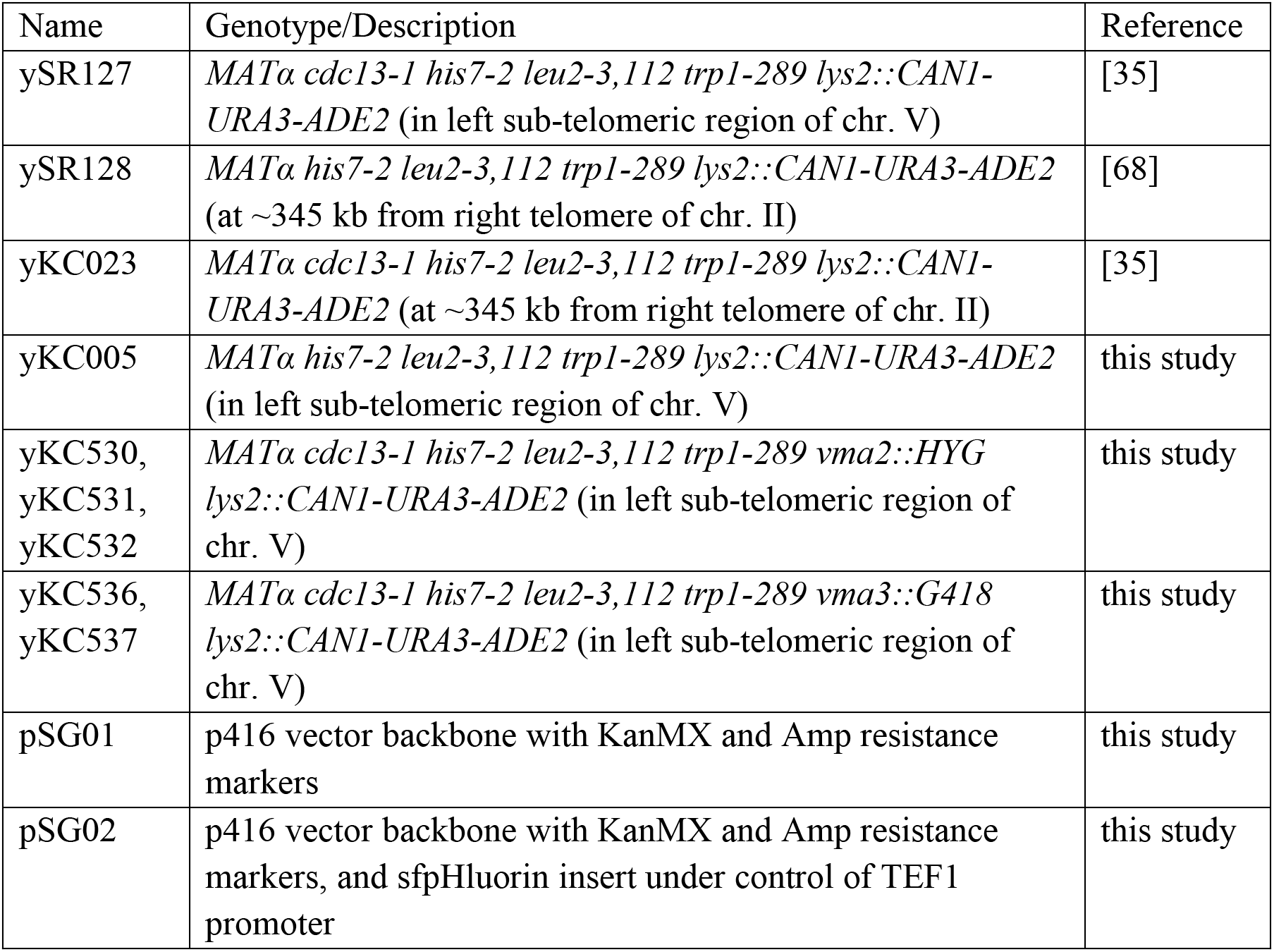
List of yeast strains and plasmids used in this work.

*vma2::HYG* and *vma3::G418* derivatives of ySR127 were constructed by one step gene replacement using the respective antibiotic resistance marker [39]. Constructions were verified by diagnostic replica plating and PCRs.

ρ^-^ petites were isolated from the ySR127 parental background as follows: Four ySR127 colonies were inoculated separately into YPDA liquid media and grown for three days at 23°C. 200 cells were plated onto YPDA solid media in five-fold replicate from each culture. Plates were incubated at 23°C for five days. ρ^-^ candidates were selected based on small colony size, patched onto fresh YPDA, and tested by replica plating onto glycerol media.

ρ^0^ petites were isolated from the ySR127 parental background as follows [40]: Overnight cultures of ySR127 in YPDA were diluted 1:10 into fresh YPDA media with either 1 or 10 μg/mL ethidium bromide, and shaken at 23°C for three hours. Afterwards, cells were washed to remove the ethidium and 100 cells were plated onto YPDA from each culture. ρ^0^ candidates were selected based on small colony size, patched onto fresh YPDA, and tested by replica plating onto glycerol media.

To construct the pSG02 plasmid, the superfolder pHluorin gene was PCR-amplified from the p426MET25_sfpHluorin (MRV55) plasmid [41] using primers that added KpnI (GGTGGTGGTACCGCAAATTAAAGCCTTCGAGC) and XbaI (GGTGGTTCTAGAATGAGCAAAGGAGAAGAACTT) restriction sites. Purified PCR product and the pSG01 plasmid were digested with these enzymes to generate compatible sticky ends. Digested PCR product and plasmid were then ligated using T4 ligase. The ligation mix was then used to transform DH5α *E. coli* competent cells. Plasmid DNA was isolated from ampicillin-resistant clones using a miniprep kit, then screened by restriction digest to identify plasmids with the correct insertion. All steps were carried out according to the respective manufacturers’ instructions.

### Mutation Frequency Assays

Mutagenesis experiments were initiated by inoculating single colonies separately into 5 mL of YPDA-based liquid rich media in round bottom glass tubes. For experiments comparing 2% vs. 8% glucose, each colony was suspended in a small volume of sterile water, then equal aliquots were inoculated into each media type. Cells were grown at 200 RPM in permissive temperature (23°C) for three days. Then, cultures were diluted 1:10 into fresh media in fresh glass tubes, shifted to restrictive temperature (37°C), and shaken gently at 150 RPM for six or 24 hours. When mutagenesis treatment was complete, cells were collected, lightly centrifuged, washed in water, and plated onto synthetic complete media to assess survival and onto canavanine-containing media with 0.33 × adenine to select for mutants (Can^R^ colonies were off-white while Can^R^ Ade^-^ colonies turned red or pink). Care was taken to handle cells very gently, as they were quite fragile after temperature shift. Further details of this plating procedure were described previously [42]. The synthetic complete and canavanine plates were incubated at 23°C for five days. Plates were then imaged by scanner and colonies were counted using the Colony Counter plugin [43] for ImageJ software [44].

### Fluctuation Assays

Mutation rates were determined using fluctuation assays adapted from [45]. Colonies were grown for 72 hours at 23°C in liquid YPDA. Then, paired 10,000-fold dilutions were made using either YPDA (2% glucose) or YPD8A (8% glucose). 200 μL from each 10,000-fold dilution was put into a well of a 96-well 2 mL deep plate. Plates were covered securely with Microseal ‘B’ film and shaken for 72 hours at 23°C. Six cultures were selected at random from each test condition for determining cell titers using a hemacytometer and appropriate dilutions were plated on synthetic complete to assess plating efficiency. The rest of the cultures were plated onto canavanine media to identify mutants. To evaluate statistical differences in mutation rates, the flan.test() function from the flan R package (version 0.9) [46] was applied, using default parameters (Luria-Delbrück distribution, maximum likelihood model). This parametric statistical test is essentially analogous to running a t-test [46].

### Extracellular pH Measurements

pH of aliquots of conditioned culture media were determined using a pH meter (model number 860031) from Sper Scientific (Scottsdale, AZ).

### Intracellular pH Measurements

A superfolder pHluorin [41] was used to measure intracellular pH. The calibration curve and the measurements were performed similarly to the protocol in [47]. Briefly, to generate the calibration curves, cultures were incubated for 24 hours in YPDA or YPD8A media, centrifuged for 5 minutes at 4000 RPM, resuspended in PBS containing 100 mg/L digitonin, and incubated at room temperature for 10 minutes. Cells then were washed with PBS and put on ice. Next, cells were suspended to obtain an OD600 of 0.5 in PBS buffer of pH values ranging from 3.5 to 9.0 and transferred to CELLSTAR black polystyrene clear bottom 96 well microtiter plates. Superfolder pHluorin fluorescence emission was measured at 506 nm using a Synergy H1 Multi-Mode Plate Reader (BioTek, Winooski, VT) providing excitation bands of 7 nm centred around 390 and 470 nm. To generate the calibration curve, we plotted the corresponding buffer pH against the ratio of 390 and 470 nm (R390/470). As a reference for background fluorescence, a parental culture (i.e., without plasmid) was always grown simultaneously with cultures carrying the pSG02 plasmid expressing the superfolder pHluorin. For the measurement of the cytosolic pH of the samples, cell suspensions were diluted to an appropriate OD600 (ranging between 0.5 to 1.0) with preconditioned media from each culture before measurements were taken at 390 and 470 nm.

### Statistical Analyses

t-tests were performed using the t.test() function in base R (version 4.1.1) [48]. Analysis of variance model fitting and Tukey Honest Significant Differences (HSD) tests were done using the aov() and TukeyHSD() functions from the stats package (version 4.1.1) of base R [48]. Additional data processing and visualizations were generated using the tidyverse packages (version 1.3.1) [49].

## Results

### Increased glucose metabolism leads to increased mutagenesis

Mutation frequency experiments were done using haploid yeast strains (the parental strain ySR127 and its derivatives) that form long stretches of sub-telomeric ssDNA when shifted to 37°C, due to the *cdc13-1* temperature-sensitive point mutation [37] (see Figure 1A). At 37°C the mutant cdc13-1 protein dissociates from telomeres, leaving the unprotected chromosome ends vulnerable to enzymatic resection. Once significant resection does occur, the DNA damage checkpoint is activated to arrest cells in G2 [37]. Three reporter genes (*CAN1, ADE2*, and *URA3*) had been deleted from their native loci and repositioned to the left sub-telomeric region of chromosome V [35]. This mutagenesis system is exceptionally well suited to investigating weak mutagens or subtle mutagenic effects, since the ssDNA is more prone to mutation than double-stranded DNA (dsDNA) and high-fidelity repair using the complementary strand would not be possible. The *cdc13-1* ssDNA triple reporter gene system has been used previously to elucidate the mutagenic properties of: sodium bisulfite and the human APOBEC3G cytidine deaminase [35]; abasic sites [50]; reactive oxygen species [51]; human APOBEC3A and APOBEC3B cytidine deaminases [52]; alkylating agents [53]; acetaldehyde [54,55]; and formaldehyde [55].

**Figure 1:**
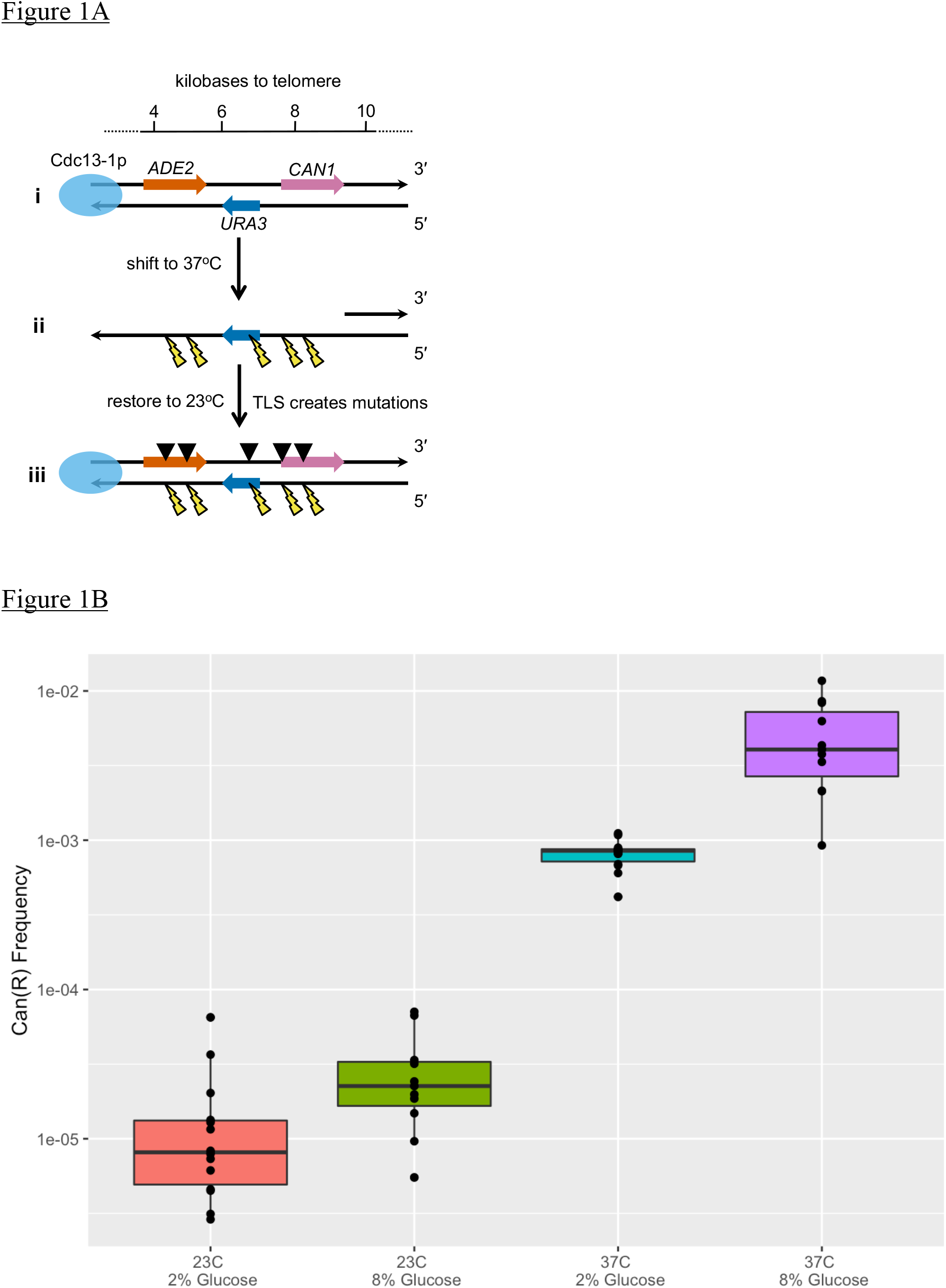
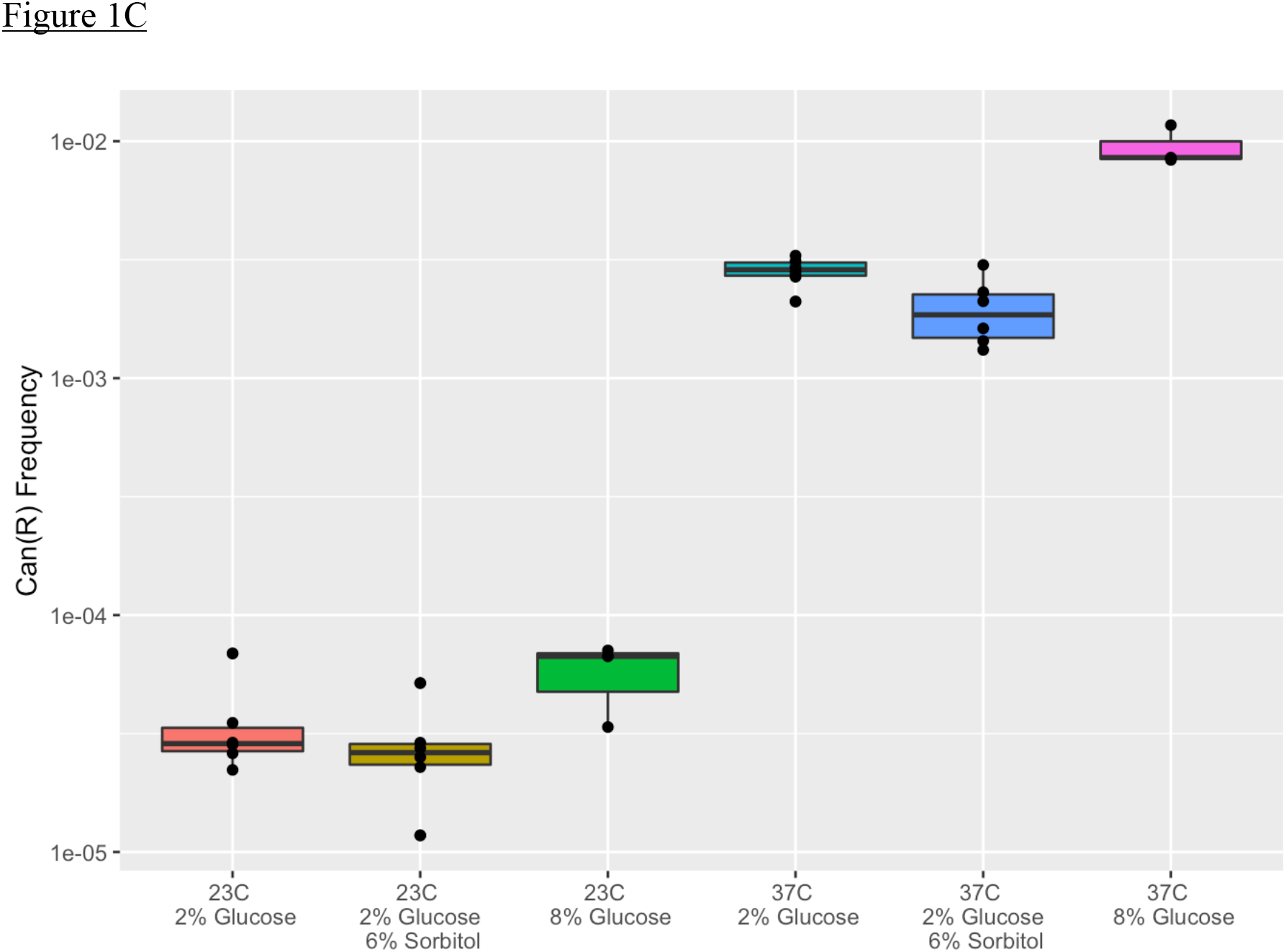
Mutagenesis depends on sugar concentration in the media. (A) Schematic diagram of *cdc13-1* sub-telomeric triple reporter gene cassette system in strain ySR127 and its derivatives. (i) Three reporter genes, *CAN1, URA3*, and *ADE2*, were relocated from their native loci to the sub-telomeric region of a de novo telomere at the end of the left arm of chromosome V. The scale at the top indicates kilobases to beginning of the nearest telomeric repeat DNA. (ii) Upon shifting to 37°C, the telomeric capping complex dissociates due to the mutant Cdc13-1 protein, exposing the chromosome end to enzymatic resection that generates long stretches of ssDNA. The long ssDNA triggers the DNA damage checkpoint to arrest cells in G2. (iii) When cells are restored to 23°C, the DNA is restored to double-stranded form by re-synthesis of the complementary strand. Error-prone translesion synthesis (TLS) polymerases can introduce mutations, which can be detected by selecting for inactivation of reporter genes. (B) Canavanine resistance (Can^R^) frequencies of ySR127 cells at 23°C or after 24-hour shift to 37°C, metabolizing media with 2% or 8% glucose. (C) Can^R^ frequencies of ySR127 cells at 23°C or after 24-hour shift to 37°C, metabolizing media with 2% glucose, 2% glucose + 6% sorbitol, or 8% glucose.

We began by comparing the mutagenic effects of metabolizing rich media with either 2% vs. 8% glucose. When ySR127 cells were grown at 23°C, there was no statistical difference in frequency of *CAN1* gene inactivation between 2% vs. 8% glucose (paired t-test P-value = 0.0858, see Figure 1B). The sub-telomeric DNA at permissive temperature should have remained mostly in the double-stranded form, so it should be better protected from mutagenesis. In contrast, when cells were shifted to 37°C for 24 hours, resulting in exposure of sub-telomeric ssDNA, there was a clearly detectable difference in mutagenesis when metabolizing 2% vs. 8% glucose (paired t-test P-value = 0.00166, see Figure 1B). These results reaffirm previous observations [34].

We then ran additional experiments to check if the excess osmolarity in the media might, in itself, affect mutagenesis. To do this, we compared mutation frequencies when metabolizing media with 2% glucose vs. 2% glucose + 6% sorbitol vs. 8% glucose. At 23°C, we did not observe a statistically significant effect on mutagenesis due to differences in media (ANOVA P-value = 0.0726, see Figure 1C). But at 37°C, we did find a significant effect on mutagenesis among the different media (ANOVA P-value = 1.74 × 10^-7^, see Figure 1C). To determine which media were statistically different from each other, we then ran the Tukey Honest Statistical Differences (HSD) test, and determined that at 37°C, the cells in 8% glucose media had higher mutation frequencies than cells in the other two media types (see Table 2). Cells metabolizing 2% glucose media did not have statistically different mutation frequencies than cells metabolizing 2% glucose + 6% sorbitol media (see Table 2). From these results, we infer that the increase in mutation frequency when metabolizing 8% glucose was not likely due to increased osmolarity in the media.

**Table 2:** Tukey HSD adjusted P-values for pairwise comparisons among media containing 2% glucose, 2% glucose + 6% sorbitol, or 8% glucose.

| Comparison | 23°C | 37°C |
| --- | --- | --- |
| 2% glucose vs. 2% glucose + 6% sorbitol | 0.743 | 0.274 |
| 2% glucose vs. 8% glucose | 0.171 | 0.0000007 |
| 2% glucose + 6% sorbitol vs. 8% glucose | 0.0620 | 0.0000002 |

We also noticed that the mutation frequency results at 23°C came close to statistical significance when comparing 2% vs. 8% glucose. To determine if longer term metabolism could reveal differences in mutagenesis, we carried out fluctuation assays to determine mutation rates at permissive temperature [56]. For ySR127, which has *cdc13-1* and the *CAN1-URA3-ADE2* reporter cassette near a chromosome end, we obtained mutation rates of 2.57 × 10^-6^ and 6.60 × 10^-6^ on 2% and 8% glucose media, respectively (flan.test parametric test P-value = 1.42 × 10^-12^), indicating a clear difference. For ySR128, which is *CDC13^+^* and has the triple reporter gene cassette in a locus in the middle of chromosome II, we found mutation rates of 1.35 × 10^-6^ on 2% glucose media and 1.80 × 10^-6^ on 8% glucose (flan.test parametric P-value = 0.0111). As an additional control, we tested strain yKC023, which has *cdc13-1*, but the *CAN1-URA3-ADE2* reporter cassette is situated at the mid-chromosome II locus. For yKC023, we determined mutation rates of 2.34 × 10^-6^ and 2.48 × 10^-6^ on 2% and 8% glucose, respectively (flan.test parametric P-value = 0.648). We also tested yKC005, which is *CDC13*^+^ but has the reporter gene cassette in the chromosome V sub-telomeric location. yKC005 yielded mutation rates of 8.93 × 10^-7^ on 2% glucose and 9.34 × 10^-7^ on 8% glucose (flan.test parametric P-value = 0.804). These data are summarized in Table 3. Consistent with the model that the SBS5-like mutagenesis in yeast is associated with ssDNA, these data showed the clearest difference in mutation rates for ySR127, which had the reporter cassette near a chromosome end and a mutation that destabilized the telomere capping complex.

**Table 3:** Summary of data from fluctuation assays to determine mutation rates of the following strains while metabolizing media with either 2% or 8% glucose: ySR127 (*cdc13-1* with sub-telomeric reporter); ySR128 (*CDC^+^* with mid-chromosome reporter); yKC023 (*cdc13-1* with mid-chromosome reporter); and yKC005 (*CDC^+^* with sub-telomeric reporter).

| Strain | Mutation Rate in<br>2% Glucose | Mutation Rate in<br>8% Glucose | flan.test Parametric<br>Test P-value |
| --- | --- | --- | --- |
| ySR127 | $2.57 \times 10^{-6}$ | $6.60 \times 10^{-6}$ | $1.42 \times 10^{-12}$ |
| ySR128 | $1.35 \times 10^{-6}$ | $1.80 \times 10^{-6}$ | 0.0111 |
| yKC023 | $2.34 \times 10^{-6}$ | $2.48 \times 10^{-6}$ | 0.648 |
| yKC005 | $8.93 \times 10^{-7}$ | $9.34 \times 10^{-7}$ | 0.804 |

### Aerobic respiration does not contribute significantly to mutagenesis

We noted previously that blocking the last step of glycolysis using a degron allele of *CDC19*, the pyruvate kinase gene expressed while metabolizing glucose [57], resulted in a ~90% decrease in mutagenesis [34]. Similarly, petite cells (lacking functional mitochondria) metabolizing 2% glucose did not exhibit any change in mutation frequency [34]. To investigate the possible role of aerobic respiration on SBS5-like mutagenesis further, we tested spontaneously occurring ρ^-^ mutants (lacking mitochondria capable of aerobic respiration) and ethidium-induced ρ^0^ mutants (lacking mitochondria altogether due to loss of mitochondrial DNA). Wild-type, ρ^-^, and ρ^0^ cells were metabolizing media with either 2% or 8% glucose, at 23°C or shifted to 37°C for six hours. There was no statistically significant effect due to mitochondrial status (ANOVA P-value = 0.187), but as expected, there was a strong effect due to temperature shifting (ANOVA P-value = 1.55 × 10^-17^, see Figure 2A). These results showed that the mutation frequencies did not depend on mitochondrial aerobic respiration, reaffirming a key role for glycolysis instead.

**Figure 2:**
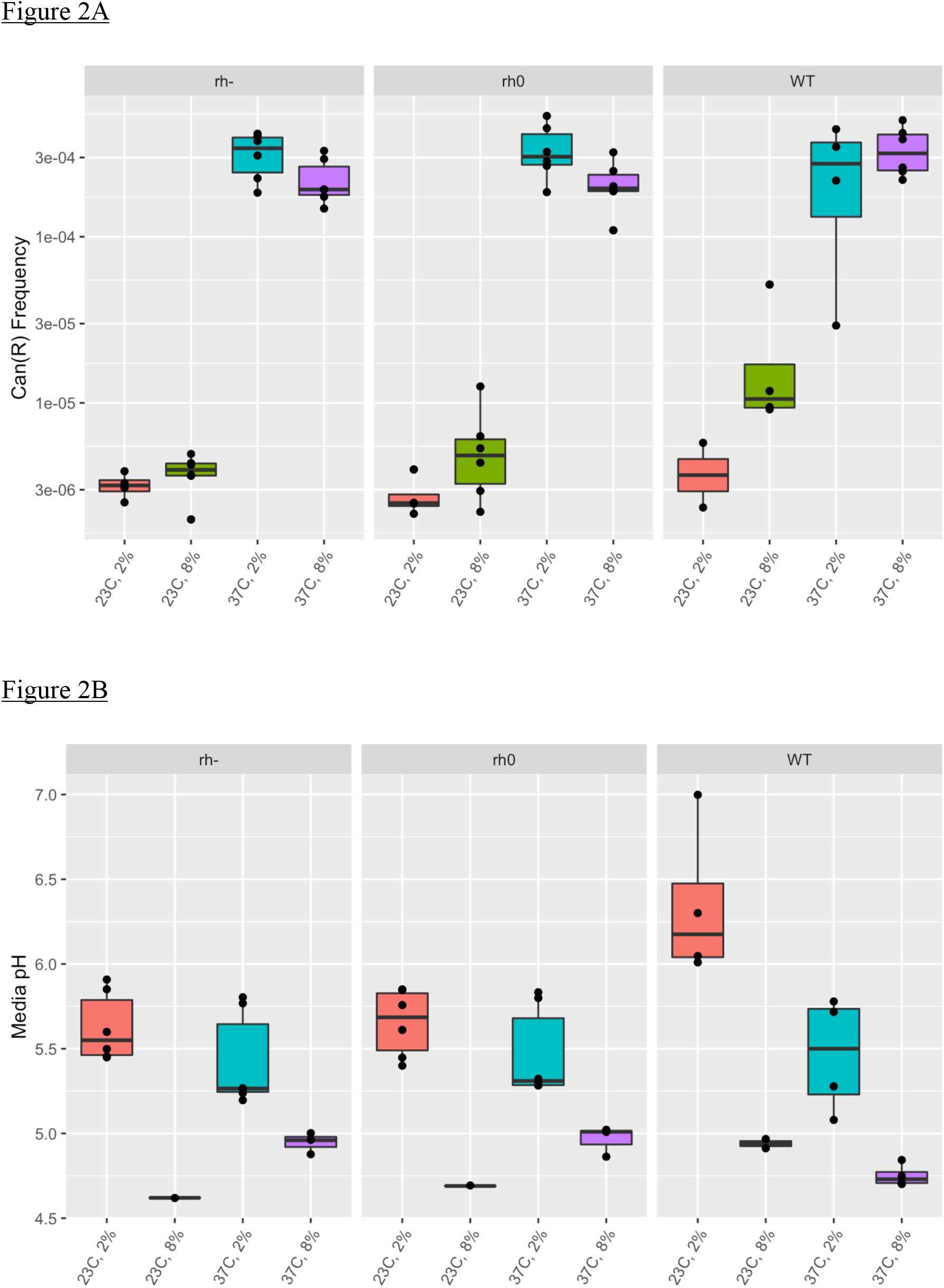
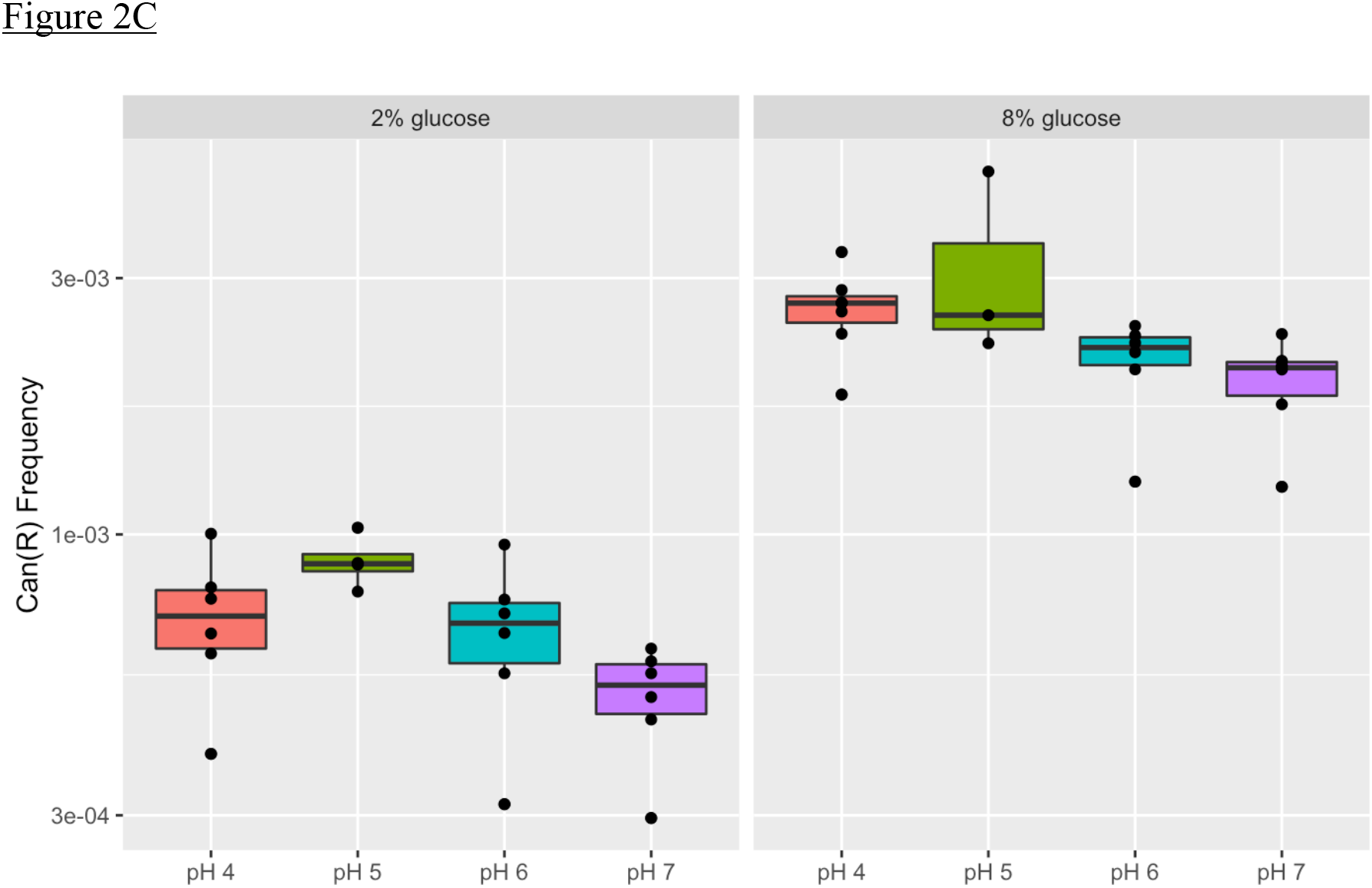
Mutagenesis does not depend on mitochondrial function, but is linked to extracellular pH. (A) Can^R^ frequencies of ρ^-^, ρ^0^, and wild-type (ySR127) cells at 23°C or after 6-hour shift to 37°C, metabolizing media with 2% or 8% glucose. (B) Media (i.e., extracellular) pH of ρ^-^, ρ^0^, and wild-type cells at 23°C or after 6-hour shift to 37°C, metabolizing media with 2% or 8% glucose. (C) Can^R^ frequencies of ySR127 cells after 24-hour shift to 37°C, metabolizing media with 2% glucose that was adjusted to the indicated pH values.

### Media acidification from metabolizing more glucose

In glycolysis, decarboxylation of pyruvate produces carbon dioxide (CO2) [58], which is rapidly hydrated by carbonic anhydrases to yield carbonic acid (H2CO3), so the net result is intracellular acidification [59]. We reasoned that if cells metabolized more glucose, then more excess protons would necessarily be produced, and more protons would have to be extruded out of the cells and into the media, as cells attempt to maintain physiological pH. This should result in a readily detectable decrease of extracellular pH in the conditioned media. We measured the pH of conditioned media for cells at 23°C or shifted for six hours to 37°C, metabolizing either 2% or 8% glucose. There was no statistical difference when comparing among the ρ status of the three genotypes (ANOVA P-value = 0.78), but there were significant differences due to treatment conditions (see Figure 2B and Table 4). There were statistically significant differences for the following pairwise comparisons of treatment conditions by the Tukey HSD test: 2% vs. 8% glucose at 23°C; 2% vs. 8% glucose at 37°C; and 23°C vs. 37°C in 2% glucose. Interestingly, there was no statistical difference for the 23°C vs. 37°C comparison in 8% glucose (Tukey HSD adjusted P-value = 0.965). Media pH was quite acidic when cells were metabolizing 8% glucose (external pH ≥ 5), regardless of other factors, i.e., either genotype or temperature. This likely reflected the necessity for all the cells to put out a higher concentration of excess protons into the media when metabolizing the higher concentration of glucose, in order to control intracellular pH.

**Table 4:** Tukey HSD adjusted P-values for pairwise comparisons of external pH among wild-type, ρ^-^, and ρ^0^ cells under different treatments.

| Comparison | Adjusted P-value |
| --- | --- |
| WT vs. $\rho^-$ | 0.761 |
| WT vs. $\rho^0$ | 0.921 |
| $\rho^-$ vs. $\rho^0$ | 0.941 |
| 23°C, 2% vs. 8% glucose | $1.0 \times 10^{-6}$ |
| 37°C, 2% vs. 8% glucose | $7.49 \times 10^{-5}$ |
| 2% glucose, 23°C vs. 37°C | $5.78 \times 10^{-3}$ |
| 8% glucose, 23°C vs. 37°C | 0.965 |

### Effect of extracellular pH on mutagenesis

We found previously that more glucose metabolism led to more mutagenesis, and have just noted that more glucose metabolism resulted in significantly more extracellular acidification. So, we next investigated if it might be possible to impinge on mutation frequency indirectly by controlling the extracellular pH. We shifted ySR127 cells to 37°C for 24 hours in either 2% or 8% glucose media that were adjusted to either pH 4, 5, 6, or 7 (see Figure 2C). ANOVA revealed a strong effect for glucose concentration on mutation frequency (P-value = 7.38 × 10^-16^), as well as a significant effect for extracellular pH (P-value = 0.00823). Using the Tukey HSD test, we found that mutation frequencies in pH 4 and 5 media both were significantly higher than in pH 7 media (see Table 5). These data show that, with a sufficiently sensitive mutation detection system, it was actually possible to detect differences in induced mutagenesis due to manipulating extracellular pH, at both glucose concentrations.

**Table 5:** Tukey HSD adjusted P-values for pairwise comparisons of mutation frequencies in cells that were temperature shifted in media with different pH.

| Comparison | Adjusted P-value |
| --- | --- |
| pH 4 vs. pH 5 | 0.918 |
| pH 4 vs. pH 6 | 0.221 |
| pH 4 vs. pH 7 | 0.0346 |
| pH 5 vs. pH 6 | 0.120 |
| pH 5 vs. pH 7 | 0.0221 |
| pH 6 vs. pH 7 | 0.825 |

### Cells metabolizing more glucose have lower intracellular pH

Given that the extracellular pH values when cells metabolize 8% glucose media were lower than cells metabolizing 2% glucose, we reasoned that it might be possible to detect differences in intracellular pH as well. To test this, we expressed a pH-sensitive superfolder pHluorin variant of GFP [41] in the ySR127 background. The distribution of intracellular pH values for cells at 23°C in 8% glucose were statistically lower than for cells in 2% glucose (paired t-test P-value = 0.00389, see Figure 3A). Similarly at 37°C after 24-hour shift, cells metabolizing 8% glucose had statistically lower intracellular pH value distributions than cells metabolizing 2% glucose (paired t-test P-value = 0.000742, see Figure 3B). These measurements show that, despite the action of homeostatic mechanisms to maintain intracellular pH within a narrow range [60], it was possible to detect differences when cells were metabolizing different concentrations of glucose glycolytically, as this metabolism would itself be the source for excess protons. This detection was made possible using a sufficiently sensitive reporter system and an experimental design with paired samples.

**Figure 3:**
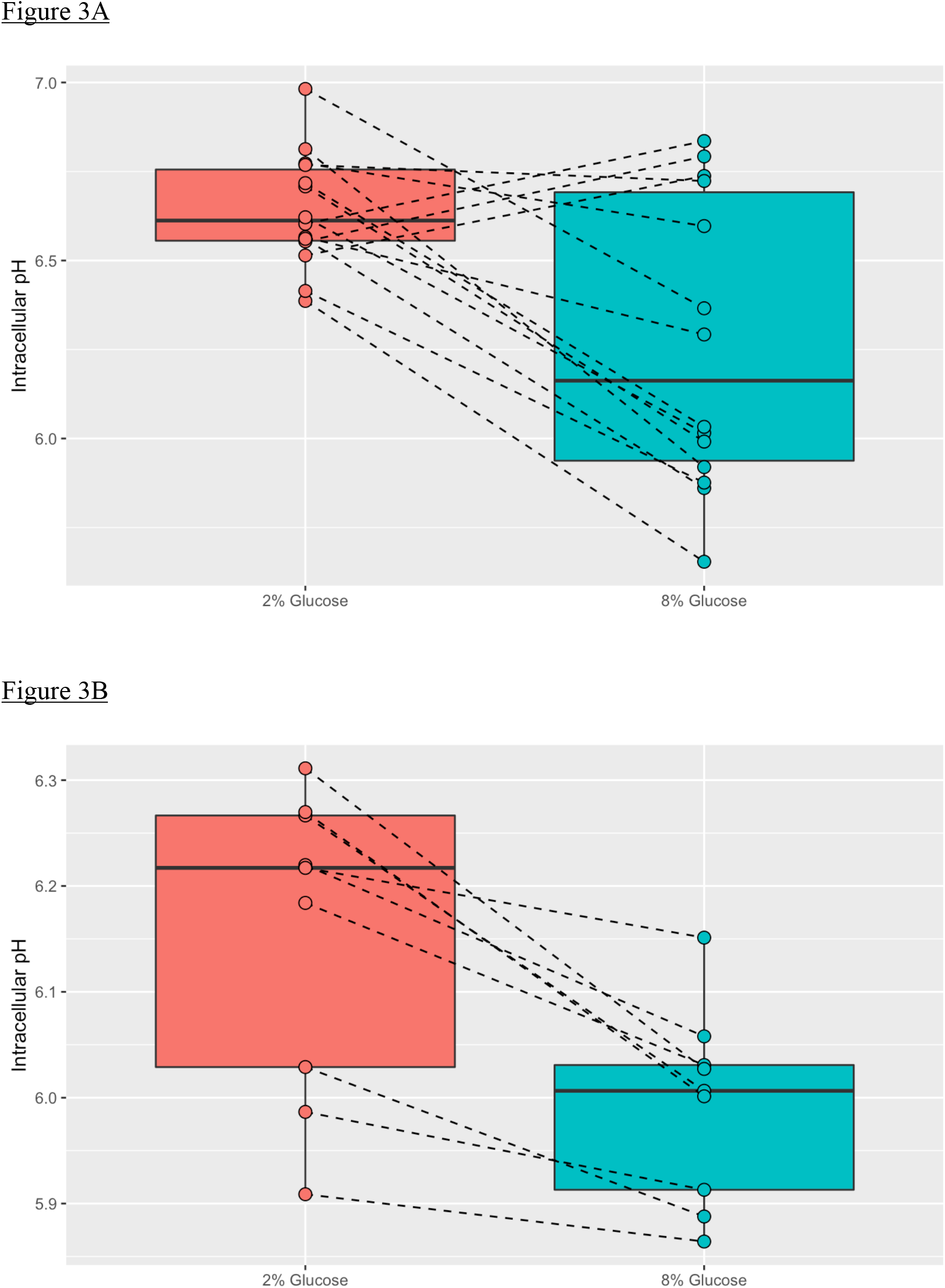
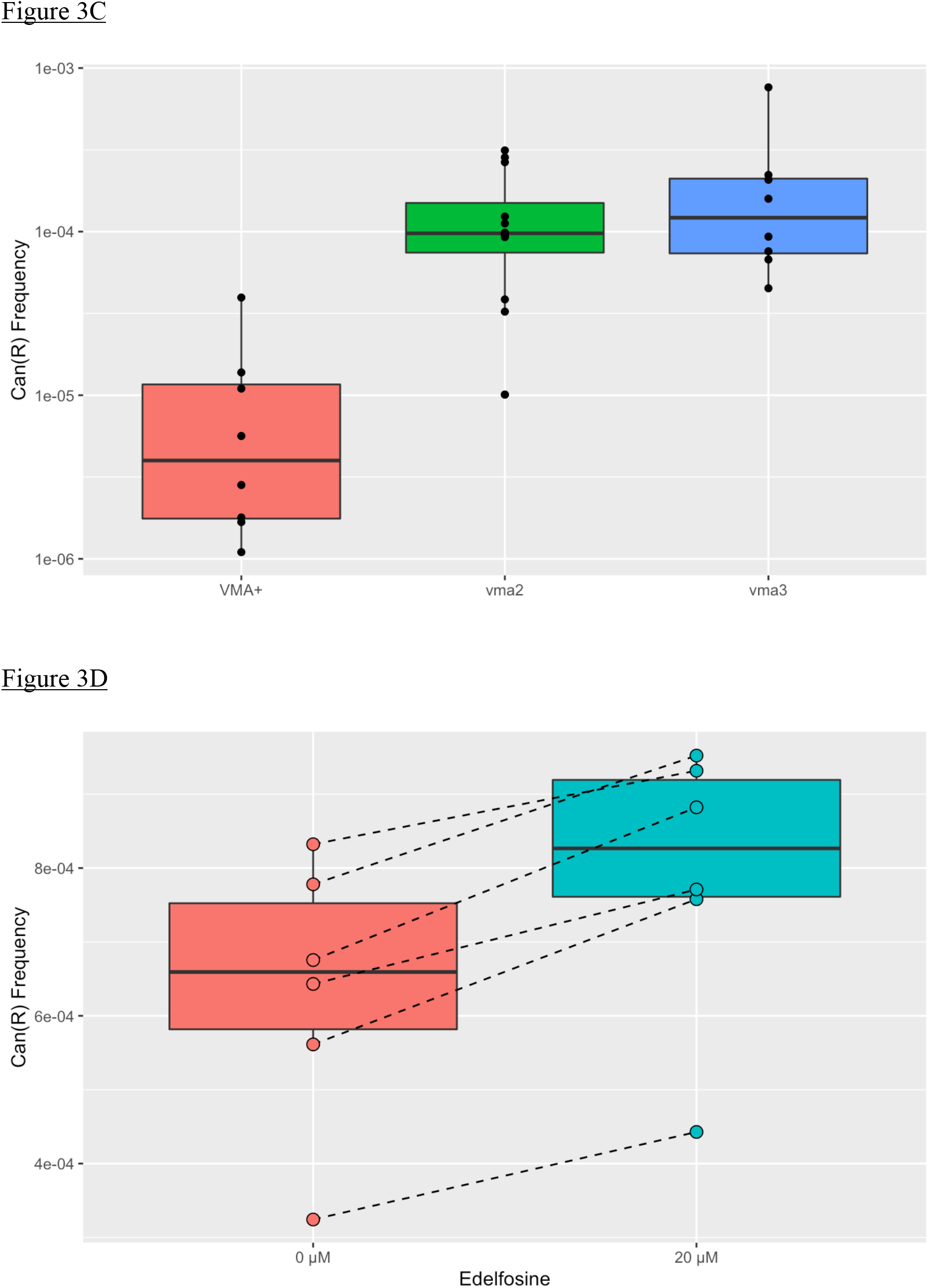
Glucose metabolism and mutagenesis are linked to intracellular pH. (A) Intracellular pH of ySR127 cells metabolizing media with 2% or 8% glucose at 23°C. (B) Intracellular pH of ySR127 cells metabolizing media with 2% or 8% glucose after 24-hour shift to 37°C. (C) Can^R^ frequencies of *vma2Δ, vma3Δ*, and wild-type (ySR127) cells at 23°C, metabolizing media with 2% glucose. (D) Can^R^ frequencies of ySR127 cells after 24-hour shift to 37°C, metabolizing media with 2% glucose and treated with either 0 or 20 μM edelfosine.

### Effect of intracellular pH on mutagenesis

Finally, we tested whether experimental perturbation of intracellular pH could affect mutation frequencies. We deleted two genes encoding subunits of the vacuolar H^+^-ATPase, namely *vma2* and *vma3* [61]. When cultured in media with 2% glucose at 23°C, ANOVA found a statistically significant difference among the three tested genotypes (P-value = 0.0182, see Figure 3C), although the Tukey HSD test showed that only the *vma3* vs. *VMA^+^* (i.e., ySR127 parental control) pairwise comparison had a statistically significant adjusted P-value (0.0137). Unfortunately, the two *vma* deletion backgrounds did not arrest properly in G2 nor survive temperature shifting to 37°C, so it was not possible to collect those data.

We also used a pharmacological approach with a membrane permeabilizing compound called edelfosine, which causes intracellular acidification [62]. Edelfosine treatment was compatible with reversible cell cycle arrest of *cdc13-1* cells. With 24-hour exposure to 20 μM edelfosine at 37°C in media with 2% glucose, we observed a statistically significant increase in mutagenesis (paired t-test P-value = 0.000378, see Figure 3D). Again, with a sufficiently sensitive experimental system, it was possible to detect differences in mutation frequency by controlling a key parameter, intracellular pH.

## Discussion

In this paper, we explored the mechanisms that contribute to intrinsic background or spontaneous mutagenesis within regions of ssDNA in a budding yeast model system, i.e., the mutagenesis observed without any added mutagen. We previously showed that the resultant mutational pattern resembles SBS signature 5 from COSMIC. Analogs of these similar mutational patterns are found in many biological systems. Moreover, the pattern of human SNPs is almost an exact match to SBS5 itself [34]. Given the ubiquitous nature of this set of related mutational patterns, we reasoned that any underlying etiology is likely to be related to some endogenous process or agent that is well conserved, and that many biological systems are exposed to routinely. We previously tested whether deficiencies in molecular detoxification of endogenously generated reactive species caused increases in mutagenesis, but did not find such connections [34]. Turning our attention to a less obvious candidate etiology, we eventually found that changing the carbon source metabolism of yeast cells did have large effects on mutagenesis within ssDNA. That set of results implicated the completion of glycolysis as an important requisite process for observing the SBS5-like mutagenesis [34]. In this study, we ran additional experiments which consolidated and reaffirmed the results showing that glucose concentration is an important determinant of mutagenesis, while mitochondrial function is dispensable, which again implicates glycolysis as a key player. Thinking about the product(s) of glycolysis that could contribute to the mutagenesis led us to investigating pH, since glycolysis is known to produce intracellular acidification [58,59].

A key factor in this study is the main experimental system that was used for most of the experiments. The *cdc13-1* ssDNA-based mutagenesis reporter system is very well suited for studying mutagenic effects that cannot be detected reliably using conventional mutagenesis reporter systems. When compared to conventional systems, where the reporter gene DNA is in double-stranded form, the ssDNA system is more sensitive to the mutagenic effects of an experimental treatment, firstly because the nitrogenous bases are simply more exposed. In addition, DNA in a persistent single-stranded configuration (as opposed to transient ssDNA from melting of the duplex form) cannot benefit from high fidelity repair mechanisms that require a complementary strand to serve as a template. Without the potential intervention by DNA repair mechanisms, the ssDNA system can provide a purer readout of the effects of mutagenesis. Depending on the mutagen, the ssDNA reporter can be two to three orders of magnitude more sensitive than dsDNA controls [35]. Indeed, this system has been deployed successfully to elucidate the characteristics of multiple mutagens already [35,50–55].

Early on in the various series of experiments, the high sensitivity of the ssDNA reporter and intracellular pH detection systems revealed to us that different yeast colonies used to grow biological replicate cultures can show considerable variance in baseline mutational frequency and pH values. To control for this apparently colony-intrinsic variance, we modified experiments to utilize paired designs whenever possible, i.e., the same colony was suspended in a small volume of water and then aliquoted equally to inoculate a pair of cultures with different experimental conditions. Using the paired experimental designs, we were able to leverage the high sensitivity of the mutagenesis and pH detection systems to best advantage.

We measured pH values when cells were metabolizing media with either 2% or 8% glucose. Determining extracellular (i.e., media) pH values was most straightforward, and showed that cells metabolizing more glucose produced more acidic conditioned media. This is as expected, since many of the excess protons from glycolysis would have to be extruded into the media, lowering its pH. We also found that the superfolder pHluorin pH-sensitive GFP reporter was sufficiently sensitive to detect a significantly lower intracellular pH when cells metabolized 8% glucose. Since pH is on a logarithmic scale, the measured difference translated to about a two-fold increase in the numbers of free protons in solution within cells. These results show that differences in pH due to different metabolic states can be measurable, in spite of the robust mechanisms for maintaining intracellular pH within a relatively narrow homeostatic range, which function specifically to counteract large shifts in pH within cells [60].

Our results also implicated pH itself as a contributing factor in mutagenesis within ssDNA. Treatment with the membrane permeabilizing drug edelfosine is known to cause intracellular acidification [62], and in our system, caused a significant increase in mutagenesis at 37°C, all while metabolizing media with 2% glucose. Similarly, deletion of *vma3* deprived cells of a mechanism for regulating intracellular pH [61], and resulted in increased mutagenesis at 23°C in 2% glucose media. Deletion of *vma2* showed a similar trend, although not statistically different vs. ySR127 (*VMA*+) controls by the Tukey HSD test. Neither *vma* mutant was able to tolerate the temperature shift to 37°C, likely because they couldn’t cope with that stressor without a functional vacuole. Nonetheless, the results did point to measurable differences in mutation frequency when intracellular pH was perturbed. In addition, the ssDNA reporter system was sufficiently sensitive to detect a small, but statistically significant change in mutation frequency when extracellular pH was varied. Tukey HSD analysis showed that the significant pairwise differences were between pH 4 vs. pH 7 and between pH 5 vs. pH 7. Another way of interpreting this result is that neutral extracellular pH had an inhibitory effect on mutagenesis in ssDNA.

We propose that excess protons from glycolysis make a plausible candidate as a ubiquitous endogenous mutagen in effect, since the protons are produced constantly as a natural by-product of glycolytic metabolism, can diffuse freely into the nucleus, and can react readily with DNA bases. The nucleus itself does not have mechanisms to regulate pH, so cytosolic acidification would also result in nuclear acidification [63]. Protonation of nitrogenous bases can change their base pairing characteristics, as has been shown for protonation of adenine, cytosine, and thymine at their most likely predicted atomic positions [22]. Biophysical experiments have also shown that protonation at N1 of adenine and N3 of cytosine were especially prevalent in ssDNA, whereas these atoms would normally be protected by base pairing in dsDNA [64]. In addition, three research groups have shown that protonation of adenine had the effect of stabilizing A^+^:C wobble base pairs especially [65–67]. This is a particularly intriguing connection, since the C > T and T > C transition mutations (which are most common in SBS5 and related mutational patterns) both can arise from the stabilized A^+^:C wobble mispairs. We propose a model whereby protonation of bases in ssDNA increases the likelihood of impeding DNA re-synthesis by a high-fidelity replicative polymerase, which in turn increases the likelihood of recruitment of the error-prone translesion DNA synthesis machinery to extend past the mispairing, resulting in increased mutation frequency. Taken altogether, this work has shed new light on intrinsic mutagenesis within ssDNA, which is similar to a common form of mutagenesis that is observed in a broad range of biological systems.

## Author Contributions

Both authors contributed to the following: Conceptualization, Data Curation, Formal Analysis, Investigation, Methodology, Validation, Visualization, and Writing - Review and Editing. S.P.G. also contributed Writing - Original Draft Preparation. K.C. additionally contributed Funding Acquisition and Supervision.

## Acknowledgments

The authors gratefully acknowledge the following funding sources, which supported this work: Natural Sciences and Engineering Research Council of Canada Grant 05973/RGPIN/2017; Ontario Early Researcher Award ER17-13-013; and Canada Research Chair 950-231842.

## Abbreviations

APOBEC: <u>apo</u>lipoprotein <u>B</u> mRNA <u>e</u>diting enzyme, <u>c</u>atalytic polypeptide
COSMIC: <u>C</u>atalog <u>o</u>f <u>S</u>omatic <u>M</u>utations <u>i</u>n <u>C</u>ancer
dsDNA: <u>d</u>ouble-<u>s</u>tranded <u>D</u>NA
HSD: <u>H</u>onest <u>S</u>tatistical <u>D</u>ifferences
NMF: <u>n</u>on-negative <u>m</u>atrix <u>f</u>actorization
SBS: <u>s</u>ingle <u>b</u>ase <u>s</u>ubstitution
SNP: <u>s</u>ingle <u>n</u>ucleotide <u>p</u>olymorphism
ssDNA: <u>s</u>ingle-<u>s</u>tranded <u>DNA</u>
TLS: <u>t</u>ranslesion <u>s</u>ynthesis
YPDA: <u>y</u>east extract, <u>p</u>eptone, (2%) <u>d</u>extrose, <u>a</u>denine media
YPD8A: <u>y</u>east extract, <u>p</u>eptone, <u>8</u>% <u>d</u>extrose, <u>a</u>denine media

## Supplemental Information Caption

<u>Supplementary Table 1</u>: Tables of all numerical values used to generate graphs depicted in the figures.

## References

[1] R. De Bont, N. van Larebeke, Endogenous DNA damage in humans: a review of quantitative data, Mutagenesis. 19 (2004) 169–185. https://doi.org/10.1093/mutage/geh025.

[2] P. Irigaray, D. Belpomme, Basic properties and molecular mechanisms of exogenous chemical carcinogens, Carcinogenesis. 31 (2010) 135–148. https://doi.org/10.1093/carcin/bgp252.

[3] H. Ikehata, T. Ono, The Mechanisms of UV Mutagenesis, J. Radiat. Res. (Tokyo). 52 (2011) 115–125. https://doi.org/10.1269/jrr.10175.

[4] D.J. Keszenman, L. Kolodiuk, J.E. Baulch, DNA damage in cells exhibiting radiation-induced genomic instability, Mutagenesis. 30 (2015) 451–458. https://doi.org/10.1093/mutage/gev006.

[5] R.R. Tice, R.B. Setlow, DNA repair and replication in aging organisms and cells, in: Finch, C.E., Schneider, E.L. (Eds.), Handb. Biol. Aging, Van Nostrand Reinhold, New York, 1985: pp. 173–224.

[6] M.K. Saparbaev, D.O. Zharkov, Glycosylase Repair, in: Ref. Module Life Sci., Elsevier, 2017. https://doi.org/10.1016/B978-0-12-809633-8.06481-5.

[7] M.S. Greenblatt, W.P. Bennett, M. Hollstein, C.C. Harris, Mutations in the p53 tumor suppressor gene: clues to cancer etiology and molecular pathogenesis, Cancer Res. 54 (1994) 4855–4878.

[8] P. Neddermann, P. Gallinari, T. Lettieri, D. Schmid, O. Truong, J. J. Hsuan, K. Wiebauer, J. Jiricny, Cloning and Expression of Human G/T Mismatch-specific Thymine-DNA Glycosylase, J. Biol. Chem. 271 (1996) 12767–12774. https://doi.org/10.1074/jbc.271.22.12767.

[9] A. Sassa, Y. Kanemaru, N. Kamoshita, M. Honma, M. Yasui, Mutagenic consequences of cytosine alterations site-specifically embedded in the human genome, Genes Environ. 38 (2016) 17. https://doi.org/10.1186/s41021-016-0045-9.

[10] B.N. Ames, M.K. Shigenaga, T.M. Hagen, Oxidants, antioxidants, and the degenerative diseases of aging, Proc. Natl. Acad. Sci. 90 (1993) 7915. https://doi.org/10.1073/pnas.90.17.7915.

[11] H.J. Helbock, K.B. Beckman, M.K. Shigenaga, P.B. Walter, A.A. Woodall, H.C. Yeo, B.N. Ames, DNA oxidation matters: the HPLC-electrochemical detection assay of 8-oxo-deoxyguanosine and 8-oxo-guanine, Proc. Natl. Acad. Sci. U. S. A. 95 (1998) 288–293.

[12] F.F. Kadlubar, K.E. Anderson, S. Häussermann, N.P. Lang, G.W. Barone, P.A. Thompson, S.L. MacLeod, M.W. Chou, M. Mikhailova, J. Plastaras, L.J. Marnett, J. Nair, I. Velic, H. Bartsch, Comparison of DNA adduct levels associated with oxidative stress in human pancreas, Mutat. Res. Mol. Mech. Mutagen. 405 (1998) 125–133. https://doi.org/10.1016/S0027-5107(98)00129-8.

[13] L.A. VanderVeen, M.F. Hashim, Y. Shyr, L.J. Marnett, Induction of frameshift and base pair substitution mutations by the major DNA adduct of the endogenous carcinogen malondialdehyde, Proc. Natl. Acad. Sci. 100 (2003) 14247. https://doi.org/10.1073/pnas.2332176100.

[14] T. Lindahl, B. Nyberg, Rate of depurination of native deoxyribonucleic acid, Biochemistry. 11 (1972) 3610–3618. https://doi.org/10.1021/bi00769a018.

[15] T. Lindahl, Instability and decay of the primary structure of DNA, Nature. 362 (1993) 709–715. https://doi.org/10.1038/362709a0.

[16] J. Nakamura, V.E. Walker, P.B. Upton, S.-Y. Chiang, Y.W. Kow, J.A. Swenberg, Highly Sensitive Apurinic/Apyrimidinic Site Assay Can Detect Spontaneous and Chemically Induced Depurination under Physiological Conditions, Cancer Res. 58 (1998) 222.

[17] J.E. Haber, DNA recombination: the replication connection, Trends Biochem. Sci. 24 (1999) 271–275. https://doi.org/10.1016/S0968-0004(99)01413-9.

[18] M.M. Vilenchik, A.G. Knudson, Endogenous DNA double-strand breaks: Production, fidelity of repair, and induction of cancer, Proc. Natl. Acad. Sci. 100 (2003) 12871. https://doi.org/10.1073/pnas.2135498100.

[19] E. Letouzé, J. Shinde, V. Renault, G. Couchy, J.-F. Blanc, E. Tubacher, Q. Bayard, D. Bacq, V. Meyer, J. Semhoun, P. Bioulac-Sage, S. Prévôt, D. Azoulay, V. Paradis, S. Imbeaud, J.-F. Deleuze, J. Zucman-Rossi, Mutational signatures reveal the dynamic interplay of risk factors and cellular processes during liver tumorigenesis, Nat. Commun. 8 (2017) 1315. https://doi.org/10.1038/s41467-017-01358-x.

[20] M. Moriya, N. Slade, B. Brdar, Z. Medverec, K. Tomic, B. Jelaković, L. Wu, S. Truong, A. Fernandes, A.P. Grollman, TP53 Mutational signature for aristolochic acid: an environmental carcinogen, Int. J. Cancer. 129 (2011) 1532–1536.https://doi.org/10.1002/ijc.26077.

[21] L.B. Alexandrov, Y.S. Ju, K. Haase, P. Van Loo, I. Martincorena, S. Nik-Zainal, Y. Totoki, A. Fujimoto, H. Nakagawa, T. Shibata, P.J. Campbell, P. Vineis, D.H. Phillips, M.R. Stratton, Mutational signatures associated with tobacco smoking in human cancer, Science. 354 (2016) 618. https://doi.org/10.1126/science.aag0299.

[22] N. Russo, M. Toscano, A. Grand, F. Jolibois, Protonation of thymine, cytosine, adenine, and guanine DNA nucleic acid bases: Theoretical investigation into the framework of density functional theory, J. Comput. Chem. 19 (1998) 989–1000. https://doi.org/10.1002/(SICI)1096-987X(19980715)19:9<989::AID-JCC1>3.0.CO;2-F.

[23] Y. Podolyan, L. Gorb, J. Leszczynski, Protonation of Nucleic Acid Bases. A Comprehensive Post-Hartree-Fock Study of the Energetics and Proton Affinities, J. Phys. Chem. A. 104 (2000) 7346–7352. https://doi.org/10.1021/jp000740u.

[24] H.R. Masoodi, S. Bagheri, M. Abareghi, The effects of tautomerization and protonation on the adenine–cytosine mismatches: a density functional theory study, J. Biomol. Struct. Dyn. 34 (2016) 1143–1155. https://doi.org/10.1080/07391102.2015.1072734.

[25] I.J. Kimsey, E.S. Szymanski, W.J. Zahurancik, A. Shakya, Y. Xue, C.-C. Chu, B. Sathyamoorthy, Z. Suo, H.M. Al-Hashimi, Dynamic basis for dG•dT misincorporation via tautomerization and ionization, Nature. 554 (2018) 195.

[26] B. Alberts, A. Johnson, J. Lewis, M. Raff, K. Roberts, P. Walter, DNA, Chromosomes and Genomes, in: Mol. Biol. Cell, 6th ed., W.W. Norton & Co., New York, 2014: pp. 173–236.

[27] L.B. Alexandrov, J. Kim, N.J. Haradhvala, M.N. Huang, A.W. Tian Ng, Y. Wu, A. Boot, K.R. Covington, D.A. Gordenin, E.N. Bergstrom, S.M.A. Islam, N. Lopez-Bigas, L.J. Klimczak, J.R. McPherson, S. Morganella, R. Sabarinathan, D.A. Wheeler, V. Mustonen, L.B. Alexandrov, E.N. Bergstrom, A. Boot, P. Boutros, K. Chan, K.R. Covington, A. Fujimoto, G. Getz, D.A. Gordenin, N.J. Haradhvala, M.N. Huang, S.M.A. Islam, M. Kazanov, J. Kim, L.J. Klimczak, N. Lopez-Bigas, M. Lawrence, I. Martincorena, J.R. McPherson, S. Morganella, V. Mustonen, H. Nakagawa, A.W. Tian Ng, P. Polak, S. Prokopec, S.A. Roberts, S.G. Rozen, R. Sabarinathan, N. Saini, T. Shibata, Y. Shiraishi, M.R. Stratton, B.T. Teh, I. Vázquez-García, D.A. Wheeler, Y. Wu, F. Yousif, W. Yu, G. Getz, S.G. Rozen, M.R. Stratton, PCAWG Mutational Signatures Working Group, PCAWG Consortium, The repertoire of mutational signatures in human cancer, Nature. 578 (2020) 94–101. https://doi.org/10.1038/s41586-020-1943-3.

[28] L.B. Alexandrov, S. Nik-Zainal, D.C. Wedge, P.J. Campbell, M.R. Stratton, Deciphering Signatures of Mutational Processes Operative in Human Cancer, Cell Rep. 3 (2013) 246–259. https://doi.org/10.1016/j.celrep.2012.12.008.

[29] COSMIC | SBS - Mutational Signatures, (n.d.). https://cancer.sanger.ac.uk/signatures/sbs/ (accessed June 2, 2022).

[30] L.B. Alexandrov, P.H. Jones, D.C. Wedge, J.E. Sale, P.J. Campbell, S. Nik-Zainal, M.R. Stratton, Clock-like mutational processes in human somatic cells, Nat. Genet. 47 (2015) 1402.

[31] Y.S. Ju, I. Martincorena, M. Gerstung, M. Petljak, L.B. Alexandrov, R. Rahbari, D.C. Wedge, H.R. Davies, M. Ramakrishna, A. Fullam, S. Martin, C. Alder, N. Patel, S. Gamble, S. O’Meara, D.D. Giri, T. Sauer, S.E. Pinder, C.A. Purdie, Å. Borg, H. Stunnenberg, M. van de Vijver, B.K.T. Tan, C. Caldas, A. Tutt, N.T. Ueno, L.J. van ’t Veer, J.W.M. Martens, C. Sotiriou, S. Knappskog, P.N. Span, S.R. Lakhani, J.E. Eyfjörd, A.-L. Børresen-Dale, A. Richardson, A.M. Thompson, A. Viari, M.E. Hurles, S. Nik-Zainal, P.J. Campbell, M.R. Stratton, Somatic mutations reveal asymmetric cellular dynamics in the early human embryo, Nature. 543 (2017) 714.

[32] R. Rahbari, A. Wuster, S.J. Lindsay, R.J. Hardwick, L.B. Alexandrov, S. Al Turki, A. Dominiczak, A. Morris, D. Porteous, B. Smith, M.R. Stratton, UK10K Consortium, M.E. Hurles, Timing, rates and spectra of human germline mutation, Nat. Genet. 48 (2015) 126.

[33] S.C. Dentro, I. Leshchiner, K. Haase, M. Tarabichi, J. Wintersinger, A.G. Deshwar, K. Yu, Y. Rubanova, G. Macintyre, J. Demeulemeester, I. Vázquez-García, K. Kleinheinz, D.G. Livitz, S. Malikic, N. Donmez, S. Sengupta, P. Anur, C. Jolly, M. Cmero, D. Rosebrock, S.E. Schumacher, Y. Fan, M. Fittall, R.M. Drews, X. Yao, T.B.K. Watkins, J. Lee, M. Schlesner, H. Zhu, D.J. Adams, N. McGranahan, C. Swanton, G. Getz, P.C. Boutros, M. Imielinski, R. Beroukhim, S.C. Sahinalp, Y. Ji, M. Peifer, I. Martincorena, F. Markowetz, V. Mustonen, K. Yuan, M. Gerstung, P.T. Spellman, W. Wang, Q.D. Morris, D.C. Wedge, P. Van Loo, S.C. Dentro, I. Leshchiner, M. Gerstung, C. Jolly, K. Haase, M. Tarabichi, J. Wintersinger, A.G. Deshwar, K. Yu, S. Gonzalez, Y. Rubanova, G. Macintyre, J. Demeulemeester, D.J. Adams, P. Anur, R. Beroukhim, P.C. Boutros, D.D. Bowtell, P.J. Campbell, S. Cao, E.L. Christie, M. Cmero, Y. Cun, K.J. Dawson, N. Donmez, R.M. Drews, R. Eils, Y. Fan, M. Fittall, D.W. Garsed, G. Getz, G. Ha, M. Imielinski, L. Jerman, Y. Ji, K. Kleinheinz, J. Lee, H. Lee-Six, D.G. Livitz, S. Malikic, F. Markowetz, I. Martincorena, T.J. Mitchell, V. Mustonen, L. Oesper, M. Peifer, M. Peto, B.J. Raphael, D. Rosebrock, S.C. Sahinalp, A. Salcedo, M. Schlesner, S.E. Schumacher, S. Sengupta, R. Shi, S.J. Shin, L.D. Stein, O. Spiro, I. Vázquez-García, S. Vembu, D.A. Wheeler, T.-P. Yang, X. Yao, K. Yuan, H. Zhu, W. Wang, Q.D. Morris, P.T. Spellman, D.C. Wedge, P. Van Loo, Characterizing genetic intra-tumor heterogeneity across 2,658 human cancer genomes, Cell. 184 (2021) 2239–2254.e39. https://doi.org/10.1016/j.cell.2021.03.009.

[34] S.P. Gelova, K.N. Doherty, S. Alasmar, K. Chan, Intrinsic base substitution patterns in diverse species reveal links to cancer and metabolism, Genetics. (2022) (accepted).

[35] K. Chan, J.F. Sterling, S.A. Roberts, A.S. Bhagwat, M.A. Resnick, D.A. Gordenin, Base Damage within Single-Strand DNA Underlies In Vivo Hypermutability Induced by a Ubiquitous Environmental Agent, PLoS Genet. 8 (2012) e1003149. https://doi.org/10.1371/journal.pgen.1003149.

[36] L.H. Hartwell, R.K. Mortimer, J. Culotti, M. Culotti, Genetic Control of the Cell Division Cycle in Yeast: V. Genetic Analysis of cdc Mutants, Genetics. 74 (1973) 267–286.

[37] B. Garvik, M. Carson, L. Hartwell, Single-stranded DNA arising at telomeres in cdc13 mutants may constitute a specific signal for the RAD9 checkpoint., Mol. Cell. Biol. 15 (1995) 6128–6138. https://doi.org/10.1128/MCB.15.11.6128.

[38] A. Morrison, J.B. Bell, T.A. Kunkel, A. Sugino, Eukaryotic DNA polymerase amino acid sequence required for 3’→ 5’ exonuclease activity., Proc. Natl. Acad. Sci. 88 (1991) 9473–9477. https://doi.org/10.1073/pnas.88.21.9473.

[39] R.J. Rothstein, One-step gene disruption in yeast, in: Methods Enzymol., Academic Press, 1983: pp. 202–211. https://doi.org/10.1016/0076-6879(83)01015-0.

[40] Goldring Elizabeth S., Grossman Lawrence I., Marmur Julius, Petite Mutation in Yeast, J. Bacteriol. 107 (1971) 377–381. https://doi.org/10.1128/jb.107.1.377-381.1971.

[41] M. Reifenrath, E. Boles, A superfolder variant of pH-sensitive pHluorin for in vivo pH measurements in the endoplasmic reticulum, Sci. Rep. 8 (2018) 11985. https://doi.org/10.1038/s41598-018-30367-z.

[42] K. Chan, Molecular Genetic Characterization of Mutagenesis Using a Highly Sensitive Single-Stranded DNA Reporter System in Budding Yeast, in: M. Muzi-Falconi, G.W. Brown (Eds.), Genome Instab. Methods Protoc., Springer New York, New York, NY, 2018: pp. 33–42. https://doi.org/10.1007/978-1-4939-7306-4_4.

[43] Colony Counter, (n.d.). https://imagej.nih.gov/ij/plugins/colony-counter.html (accessed June 18, 2022).

[44] C.A. Schneider, W.S. Rasband, K.W. Eliceiri, NIH Image to ImageJ: 25 years of image analysis, Nat. Methods. 9 (2012) 671–675.

[45] Measuring Microbial Mutation Rates with the Fluctuation Assay., United States, 2019. https://doi.org/10.3791/60406.

[46] A. Mazoyer, R. Drouilhet, S. Despréaux, B. Ycart, flan: An R Package for Inference on Mutation Models, R J. 1 (2017) 334–351.

[47] R. Orij, J. Postmus, A. Ter Beek, S. Brul, G.J. Smits, In vivo measurement of cytosolic and mitochondrial pH using a pH-sensitive GFP derivative in Saccharomyces cerevisiae reveals a relation between intracellular pH and growth, Microbiology,. 155 (2009) 268–278.

[48] R Core Team, R: The R Project for Statistical Computing, (2020). https://www.r-project.org/ (accessed March 11, 2020).

[49] H. Wickham, M. Averick, J. Bryan, W. Chang, L. McGowan, R. François, G. Grolemund, A. Hayes, L. Henry, J. Hester, M. Kuhn, T. Pedersen, E. Miller, S. Bache, K. Müller, J. Ooms, D. Robinson, D. Seidel, V. Spinu, K. Takahashi, D. Vaughan, C. Wilke, K. Woo, H. Yutani, Welcome to the tidyverse, J Open Source Softw. 4 (2019) 1686. https://doi.org/10.21105/joss.01686.

[50] K. Chan, M.A. Resnick, D.A. Gordenin, The choice of nucleotide inserted opposite abasic sites formed within chromosomal DNA reveals the polymerase activities participating in translesion DNA synthesis, DNA Repair. 12 (2013) 878–889. https://doi.org/10.1016/j.dnarep.2013.07.008.

[51] N.P. Degtyareva, L. Heyburn, J. Sterling, M.A. Resnick, D.A. Gordenin, P.W. Doetsch, Oxidative stress-induced mutagenesis in single-strand DNA occurs primarily at cytosines and is DNA polymerase zeta-dependent only for adenines and guanines, Nucleic Acids Res. 41 (2013) 8995–9005. https://doi.org/10.1093/nar/gkt671.

[52] K. Chan, S.A. Roberts, L.J. Klimczak, J.F. Sterling, N. Saini, E.P. Malc, J. Kim, D.J. Kwiatkowski, D.C. Fargo, P.A. Mieczkowski, G. Getz, D.A. Gordenin, An APOBEC3A hypermutation signature is distinguishable from the signature of background mutagenesis by APOBEC3B in human cancers, Nat Genet. 47 (2015) 1067–1072.

[53] N. Saini, J.F. Sterling, C.J. Sakofsky, C.K. Giacobone, L.J. Klimczak, A.B. Burkholder, E.P. Malc, P.A. Mieczkowski, D.A. Gordenin, Mutation signatures specific to DNA alkylating agents in yeast and cancers, Nucleic Acids Res. 48 (2020) 3692–3707. https://doi.org/10.1093/nar/gkaa150.

[54] S. Vijayraghavan, L. Porcher, P.A. Mieczkowski, N. Saini, Acetaldehyde makes a distinct mutation signature in single-stranded DNA., Nucleic Acids Res. 50 (2022) 7451–7464. https://doi.org/10.1093/nar/gkac570.

[55] M.J. Thapa, R.M. Fabros, S. Alasmar, K. Chan, Analyses of Mutational Patterns Induced by Formaldehyde and Acetaldehyde Reveal Similarity to a Common Mutational Signature, G3 GenesGenomesGenetics. (2022) jkac238. https://doi.org/10.1093/g3journal/jkac238.

[56] S.A. Nick McElhinny, D.A. Gordenin, C.M. Stith, P.M.J. Burgers, T.A. Kunkel, Division of Labor at the Eukaryotic Replication Fork, Mol. Cell. 30 (2008) 137–144. https://doi.org/10.1016/j.molcel.2008.02.022.

[57] R.L. Burke, P. Tekamp-Olson, R. Najarian, The isolation, characterization, and sequence of the pyruvate kinase gene of Saccharomyces cerevisiae., J. Biol. Chem. 258 (1983) 2193–2201.

[58] J.T. Pronk, H. Yde Steensma, J.P. Van Dijken, Pyruvate Metabolism in Saccharomyces cerevisiae, Yeast. 12 (1996) 1607–1633. https://doi.org/10.1002/(SICI)1097-0061(199612)12:16<1607::AID-YEA70>3.0.CO;2-4.

[59] B.C. Tripp, K. Smith, J.G. Ferry, Carbonic Anhydrase: New Insights for an Ancient Enzyme, J. Biol. Chem. 276 (2001) 48615–48618. https://doi.org/10.1074/jbc.R100045200.

[60] R. Orij, S. Brul, G.J. Smits, Intracellular pH is a tightly controlled signal in yeast, Syst. Biol. Microorg. 1810 (2011) 933–944. https://doi.org/10.1016/j.bbagen.2011.03.011.

[61] M. Forgac, Structure and Properties of the Vacuolar (H+)-ATPases *, J. Biol. Chem. 274 (1999) 12951–12954. https://doi.org/10.1074/jbc.274.19.12951.

[62] O. Czyz, T. Bitew, A. Cuesta-Marbán, C.R. McMaster, F. Mollinedo, V. Zaremberg, Alteration of Plasma Membrane Organization by an Anticancer Lysophosphatidylcholine Analogue Induces Intracellular Acidification and Internalization of Plasma Membrane Transporters in Yeast, J. Biol. Chem. 288 (2013) 8419–8432. https://doi.org/10.1074/jbc.M112.425744.

[63] J.R. Casey, S. Grinstein, J. Orlowski, Sensors and regulators of intracellular pH, Nat. Rev. Mol. Cell Biol. 11 (2010) 50–61. https://doi.org/10.1038/nrm2820.

[64] M.C. Williams, J.R. Wenner, I. Rouzina, V.A. Bloomfield, Effect of pH on the Overstretching Transition of Double-Stranded DNA: Evidence of Force-Induced DNA Melting, Biophys. J. 80 (2001) 874–881. https://doi.org/10.1016/S0006-3495(01)76066-3.

[65] T. Brown, G.A. Leonard, E.D. Booth, G. Kneale, Influence of pH on the conformation and stability of mismatch base-pairs in DNA, J. Mol. Biol. 212 (1990) 437–440. https://doi.org/10.1016/0022-2836(90)90320-L.

[66] H.T. Allawi, J. SantaLucia, Nearest-Neighbor Thermodynamics of Internal A·C Mismatches in DNA: Sequence Dependence and pH Effects, Biochemistry. 37 (1998) 9435–9444. https://doi.org/10.1021/bi9803729.

[67] N.A. Siegfried, B. O’Hare, P.C. Bevilacqua, Driving Forces for Nucleic Acid pKa Shifting in an A+·C Wobble: Effects of Helix Position, Temperature, and Ionic Strength, Biochemistry. 49 (2010) 3225–3236. https://doi.org/10.1021/bi901920g.

[68] S.A. Roberts, J. Sterling, C. Thompson, S. Harris, D. Mav, R. Shah, L.J. Klimczak, G.V. Kryukov, E. Malc, P.A. Mieczkowski, M.A. Resnick, D.A. Gordenin, Clustered Mutations in Yeast and in Human Cancers Can Arise from Damaged Long Single-Strand DNA Regions, Mol. Cell. 46 (2012) 424–435. https://doi.org/10.1016/j.molcel.2012.03.030.

